# Derivation and theoretical validation of fractional quasi-steady state approximation (fQSSA) for target-mediated drug disposition models with memory effects

**DOI:** 10.64898/2026.04.27.721025

**Authors:** Jong Hyuk Byun, Issac Park, Hwi-yeol Yun, Jae Kyoung Kim

**Affiliations:** Department of Mathematics and Institute of Mathematical Science, Pusan National University, Busan, Republic of Korea; Institute of Future Earth, Pusan National University, Busan, Republic of Korea; College of Pharmacy, Chungnam National University, Daejeon, Republic of Korea; Department of Bio-AI Convergence, Chungnam National University, Daejeon, Republic of Korea; Biomedical Mathematics Group, Institute for Basic Science, Daejeon, Republic of Korea; Department of Mathematical Sciences, KAIST, Daejeon, Republic of Korea; Department of Medicine, Korea University, Seoul, Republic of Korea

## Abstract

Standard target-mediated drug disposition (TMDD) models are widely used to describe nonlinear pharmacokinetics driven by high-affinity drug–target interactions. However, their reliance on instantaneous binding limits their ability to capture delayed and history-dependent dynamics observed *in vivo*. Here, we introduce a fractional TMDD model that incorporates memory effects through a fractional derivative, thereby generalizing the standard TMDD (sTMDD) framework. Although this fractional TMDD (fTMDD) formulation increases modeling flexibility, it also exacerbates parameter identifiability challenges under typical experimental conditions where only drug concentration data are available. To address this limitation, we derive a fractional quasi-steady-state approximation (fQSSA) that reduces model dimensionality while preserving essential nonlinear and memory-dependent pharmacokinetic dynamics. We further establish an explicit validity condition that quantifies the approximation error of both fTMDD and fQSSA without requiring numerical simulation. This condition reveals that the initial drug-to-target ratio is the primary determinant of QSSA validity, whereas the fractional order has a comparatively minor influence. Application of the proposed framework to recombinant human erythropoietin (rhEPO) data demonstrates that fractional dynamics play a population-dependent role, improving model performance in adults but not in infants. Together, this work provides the first systematic derivation of a QSSA framework for fractional TMDD models, along with rigorous and computable applicability conditions. Our results establish a principled foundation for incorporating memory effects into pharmacokinetic modeling and offer a generalizable framework for nonlinear PK–PD systems involving binding-mediated dynamics.

**Author summary:** Many drugs interact strongly with their biological targets, leading to complex and nonlinear pharmacokinetics that are commonly described using target-mediated drug disposition (TMDD) models. However, these models assume that drug–target interactions occur instantaneously, which limits their ability to capture delayed and history-dependent behaviors observed in real biological systems. In this study, we develop a new modeling framework that incorporates such memory effects by extending TMDD models using fractional calculus. To make the model more practical and computationally efficient, we derive a simplified version based on a quasi-steady-state approximation (QSSA) and provide a clear mathematical condition that determines when this simplification is valid. Our analysis shows that the accuracy of the simplified model is primarily controlled by the initial ratio of drug to target, while the influence of memory effects is comparatively smaller. When applied to experimental data for erythropoietin, our model reveals that memory effects are important in adults but negligible in infants, suggesting that these effects may reflect underlying physiological differences. Overall, this work provides a systematic and interpretable framework for incorporating memory effects into pharmacokinetic modeling, with potential applications to a wide range of drug systems involving complex binding dynamics.

## Introduction

Target-mediated drug disposition (TMDD) describes the nonlinear pharmacokinetics of drugs whose disposition is substantially influenced by high-affinity binding to their biological targets [1, 2]. Such nonlinearity arises from binding, internalization, and degradation of drug-target complexes, often resulting in saturation behavior, especially at low doses. TMDD is widely observed in biologics, including monoclonal antibodies and antibody–drug conjugates, where drug disposition is closely coupled to target expression and turnover [3–7]. Similar nonlinear pharmacokinetics have also been reported for small-molecule drugs [8, 9]. Accurate TMDD modeling is therefore important for dose selection and for predicting therapeutic efficacy and safety [2, 6, 8].

The standard TMDD (sTMDD) model, based on mass-action kinetics, characterizes the system using three primary state variables: free drug concentration (*C*), free target concentration (*R*), and drug–target complex (*RC*) (Fig 1a) [10, 11]. In more complex physiological settings, such as in the presence of drug–drug interactions, TMDD dynamics can be further modulated by competitive or uncompetitive mechanisms, leading to altered pharmacokinetic behavior [12–15]. Despite its widespread use, the sTMDD framework assumes instantaneous mass-action kinetics, meaning that binding depends only on the current drug concentration. This assumption neglects physiological memory effects, including delayed conformational changes, slow receptor dynamics, and history-dependent drug responses [16–20].

**Fig 1.**
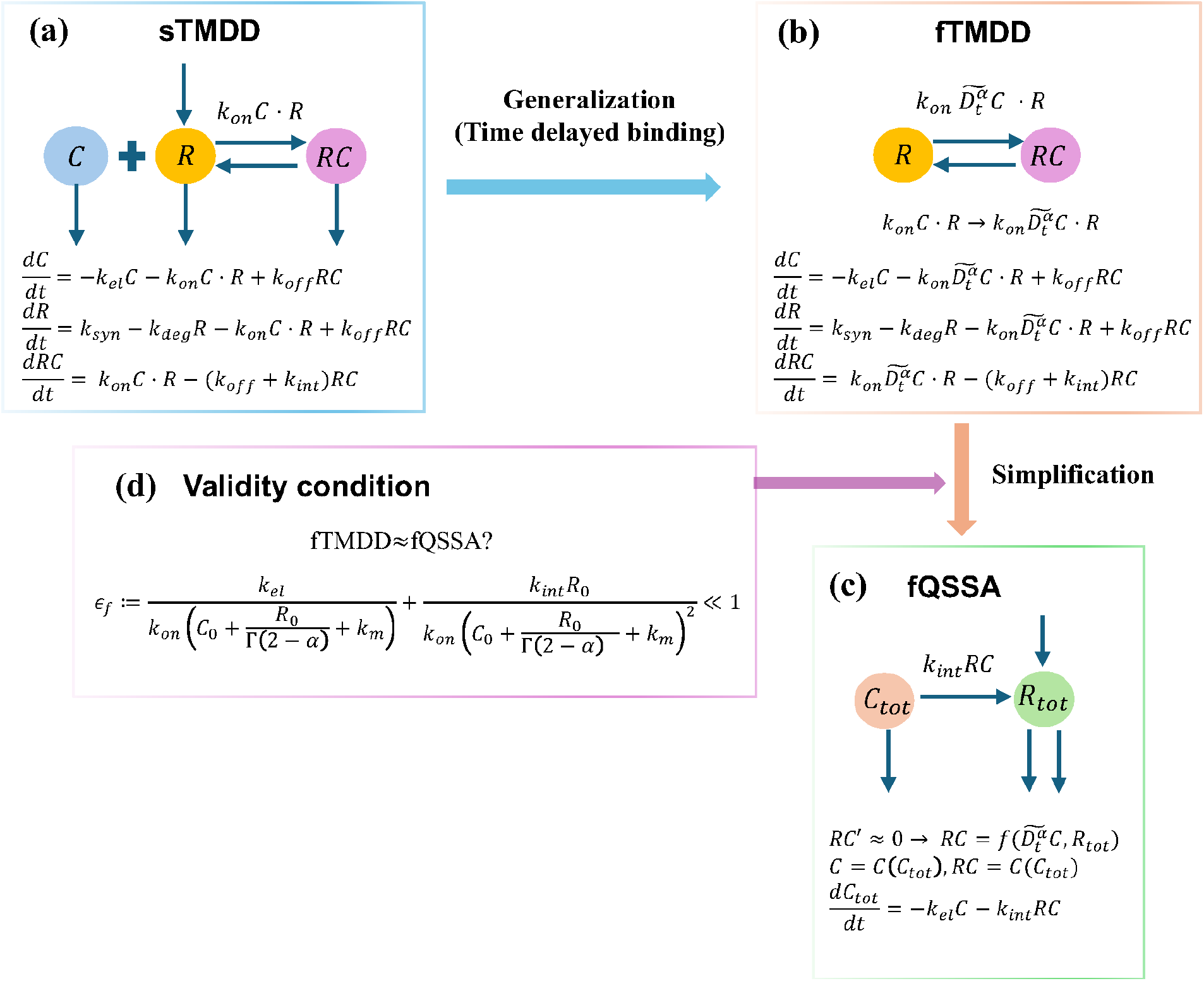
Schematic diagram of the proposed modeling framework. **(a)** The sTMDD model based on mass-action binding kinetics, describing the dynamics of the drug (*C*), target (*R*), and drug–target complex (*RC*) using a system of ODEs. **(b)** The fTMDD model, obtained as a continuous generalization of the sTMDD by applying a fractional derivative to the drug concentration *C* to account for drug conformational dynamics and delayed activation in drug-target interactions. **(c)** The fQSSA model, derived by exploiting time-scale separation between fast complex formation and slower drug dynamics, leading to an algebraic approximation for the rapidly varying complex *RC*. **(d)** Validity condition providing an *a priori* criterion to determine when the fQSSA accurately approximates the fTMDD model, without requiring numerical simulation.

To account for such delays, pharmacokinetic and pharmacodynamic (PK–PD) models have employed kernel-based or delay formulations that capture non-instantaneous effects (Fig 2, left) [21, 22]. However, these approaches typically impose memory effects from the initial time point, which can distort early-time dynamics and obscure the interpretation of initial conditions, particularly when delays emerge progressively rather than instantaneously. In contrast, fractional calculus provides a principled framework for describing history-dependent dynamics through power-law memory kernels (Fig 2, right) [23–25]. While fractional models have been successfully applied to drug absorption and elimination processes, their systematic integration into target-mediated drug disposition (TMDD) models—together with model reduction and rigorous validity analysis—remains largely unexplored.

**Fig 2.**
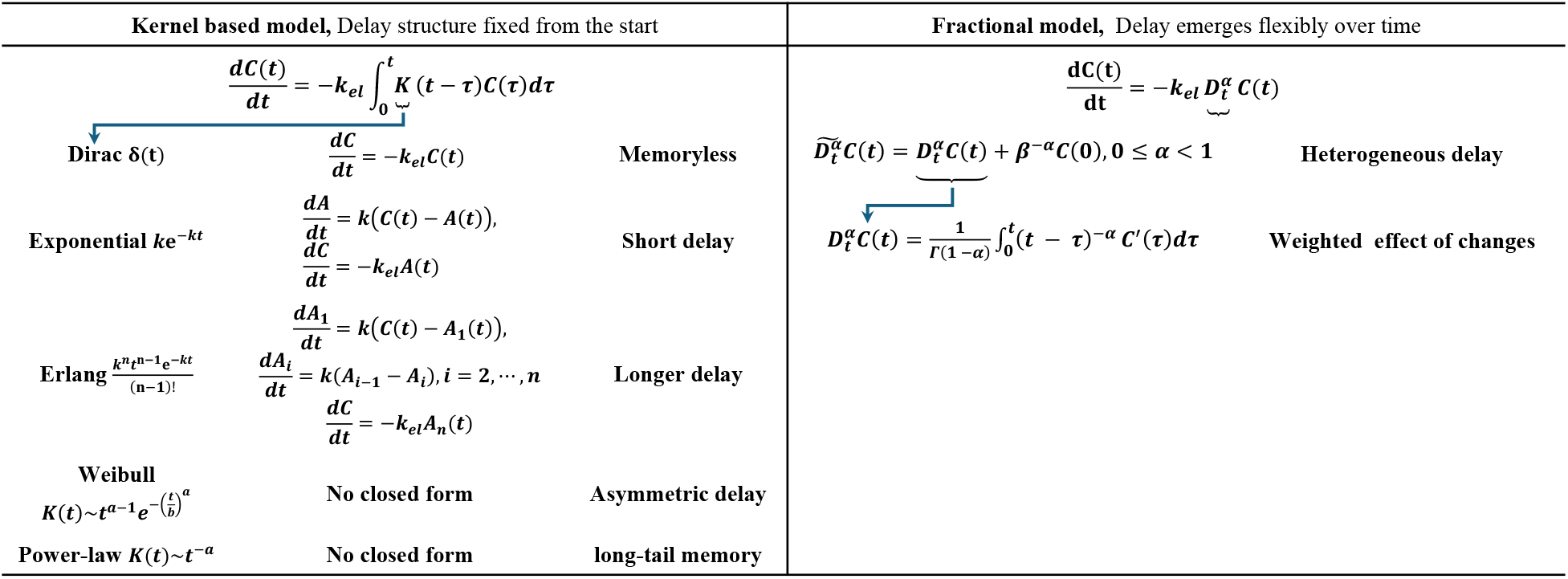
Comparison of delay structures in kernel-based and fractional models. Kernel-based models prescribe delay characteristics explicitly from the initial time through predefined kernels, whereas fractional models generate effective delays dynamically through history-dependent dynamics.

In this study, we introduce a fractional-order TMDD (fTMDD) model by incorporating a Caputo fractional derivative into the drug dynamics, enabling history-dependent binding kinetics while recovering the standard TMDD framework in the memory-free limit. Although the fTMDD improves flexibility in capturing memory effects, it increases model complexity and raises identifiability challenges, particularly under sparse data. To address this, we derive a fractional quasi-steady-state approximation (fQSSA) that reduces model dimensionality while preserving essential nonlinear dynamics. We further establish an explicit validity condition that quantifies approximation accuracy without requiring numerical simulation. Our analysis shows that the initial drug-to-target ratio primarily governs the validity of the QSSA, whereas the contribution of memory effects is comparatively minor. Application to recombinant human erythropoietin (rhEPO) data reveals a population-dependent role of memory, providing biological evidence that fractional dynamics are relevant in adults but negligible in infants. To the best of our knowledge, this work provides the first systematic derivation of a QSSA framework for fractional TMDD models together with rigorous applicability conditions. More broadly, it establishes a principled foundation for incorporating memory effects into pharmacokinetic modeling and for extending these ideas to nonlinear PK–PD systems.

## Materials and Methods

### Description of sTMDD model

The sTMDD model assumes a well-stirred, single-compartment system with intravenous (IV) bolus administration, in which the free target concentration *R* (e.g., unit nM) is synthesized at a constant rate *k*_*syn*_ (nM · h^−1^), h representing hours or days, and degraded via a first-order process with rate constant *k*_*deg*_ (1/h) [10, 11, 26]. The system is described by the following ODEs (Fig 1a):

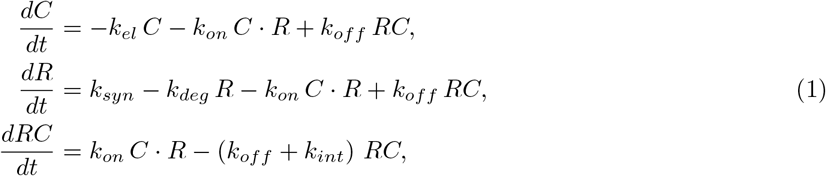

In the model, the free drug *C* (nM) binds to the target with association and dissociation rate constants *k*_*on*_ (nM^−1^*/*h) and *k*_*off*_ (1/h), respectively, while drug-target complex *RC* is internalized at rate *k*_*int*_ (1/h) and the free drug *C* is cleared via first-order elimination with rate constant *k*_*el*_ (1/h). The initial conditions are *C*(0) = *C*_0_, *R*(0) = *k*_*syn*_*/k*_*deg*_, and *RC*(0) = 0, where *R*(0) := *R*_0_ = *k*_*syn*_*/k*_*deg*_ corresponds to the pre-dose steady-state level of the free target.

### Derivation of the fTMDD model

Delayed drug conformational changes, slow redistribution between binding-competent states, and other history-dependent mechanisms can introduce memory effects that are not captured by instantaneous mass-action kinetics. To account for these effects in the sTMDD model, we introduce a fractional-order representation of the drug concentration in the drug–target binding process. Specifically, we replace the instantaneous concentration *C*(*t*) in the binding term with a modified Caputo fractional derivative:

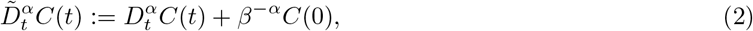

where 0 ≤ *α <* 1, Γ(·) denotes the Gamma function, and the Caputo derivative is defined as

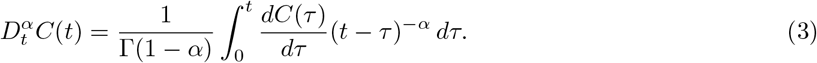

The additional term *β*^−*α*^*C*(0) ensures recovery of the standard TMDD model in the limit *α* → 0 [27]. The parameter *β* controls the decay of the memory associated with the initial condition, with smaller values corresponding to more persistent effects. In this study, we fix *β* = 1 without loss of generality. Biologically, *α* therefore quantifies the intensity of history dependence in the drug–target interaction.

Using the modified fractional derivative (Eq. 2), the fTMDD model (Fig 1b) is given by

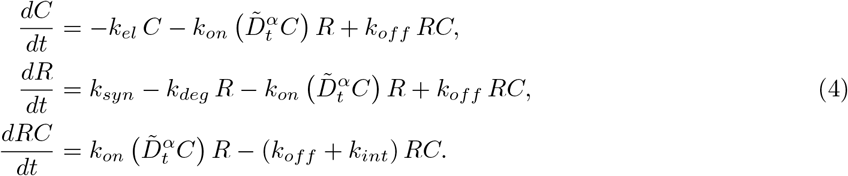

This formulation yields a hybrid FDE–ODE system in which memory effects enter exclusively through the drug concentration. Restricting the fractional derivative to *C*(*t*) is both biologically and mathematically well motivated. Biologically, delayed dynamics in drug–receptor systems primarily reflect drug-side processes rather than target turnover. Mathematically, the Caputo derivative does not satisfy a general product rule [23], rendering expressions such as 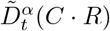 ambiguous and difficult to interpret. The present formulation therefore preserves consistency with the sTMDD framework while enabling a systematic description of history-dependent drug–target interactions.

### Derivation of the fQSSA model

To reduce the complexity of the fTMDD model, we exploit the separation of time scales and derive a fractional quasi-steady-state approximation (fQSSA). Assuming that complex formation occurs on a fast time scale (i.e., 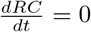 in Eq. 4), we get

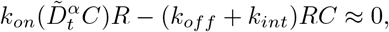

which yields

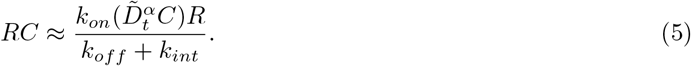

Defining *R*_*tot*_ = *R* + *RC* and *k*_*m*_ = (*k*_*off*_ + *k*_*int*_)*/k*_*on*_, we obtain

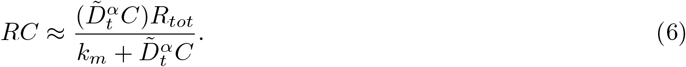

The total drug concentration *C*_*tot*_ = *C* + *RC* satisfies

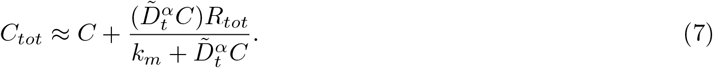

Eq. 7 is implicit in the free drug concentration *C*(*t*) because the fractional derivative 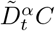 depends on the entire history of *C*. Consequently, *C*(*t*) cannot be expressed in closed form in terms of *C*_*tot*_ and must be recovered numerically. Thus, at each time point, we solve the nonlinear equation

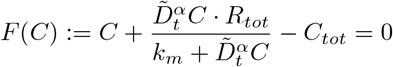

using a Newton-type method (MATLAB `fsolve` with a nonnegative constraint on the solution.). This step reconstructs the free drug concentration *C*(*t*) from *C*_*tot*_(*t*) while preserving the full memory dependence of the model.

Once *C*(*t*) is obtained, the remaining variables follow from

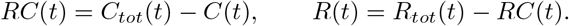

Summing the equations for *C* and *RC*, and for *R* and *RC* in Eq. 4, we obtain the fQSSA system for the total concentrations (Fig 1c):

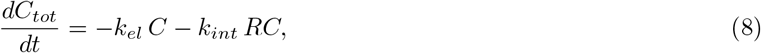

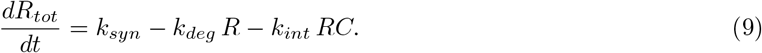

Here, *C, R*, and *RC* are implicitly defined through Eqs. 6–7 and the numerical reconstruction described above. The resulting fQSSA model reduces the dimensionality of the original fTMDD system while preserving both nonlinear and memory-dependent dynamics.

### Validity condition of the fTMDD model

While the fQSSA model provides a computationally efficient reduction of the fTMDD system, its accuracy depends on whether the drug–target complex *RC* relaxes to a quasi-steady state on a sufficiently fast timescale in the presence of memory effects. We therefore derive an explicit validity condition in terms of model parameters and initial concentrations, without relying on numerical simulation.

The fQSSA is valid if *RC* approaches its quasi-steady state within a short transient interval *t*_*c*_, during which the relative change in the total drug concentration *C*_*tot*_ remains small. Biologically, this requirement ensures that complex formation equilibrates rapidly compared to the overall pharmacokinetic dynamics. To formalize this condition, we estimate both the characteristic transient timescale *t*_*c*_ and an upper bound for *RC*(*t*) over *t* ≤ *t*_*c*_.

For 0 ≤ *α <* 1 and sufficiently small *t*, we have *C*_*tot*_(*t*) ≈ *C*_0_ and *R*_*tot*_(*t*) ≈ *R*_0_, so that *C*(*t*) ≈ *C*_0_ − *RC*(*t*). Taking the Caputo derivative yields

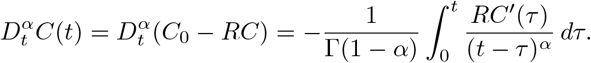

During the initial binding transient *RC*(*t*) increases monotonically from zero, and a first-order scaling *RC*^*′*^(*τ*) = *RC*(*t*)*/t, τ* ∈ (0, *t*) provides a conservative bound by mean value theorem. This leads to the approximation

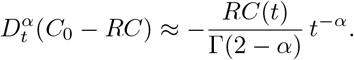

Using the definition of the modified Caputo operator, we obtain

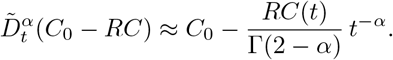

Substituting this expression into the *RC* equation in Eq. 4, imposing *RC*(0) = 0, and neglecting higher-order terms in *RC*, we obtain

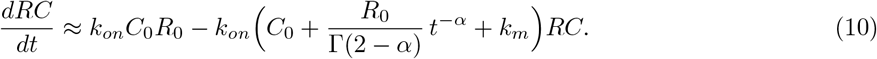

For *t* ∈ (0, *t*_*c*_], we have *t*^−*α*^ ≥ 1, which yields the differential inequality

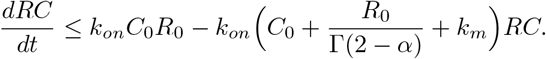

Solving this inequality gives

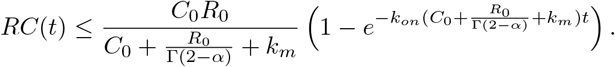

The characteristic transient timescale of complex formation is therefore estimated as

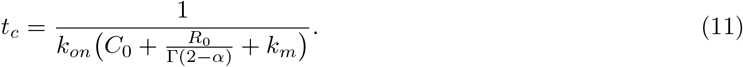

Using the mean value theorem and the fact that *t*_*c*_ is small, we obtain the simplified bound

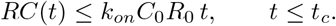

The relative change in the total drug concentration over the transient interval [0, *t*_*c*_] can now be bounded as

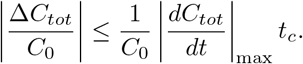

Approximating *dC*_*tot*_*/dt* using *C*_*tot*_ ≈ *C*_0_ during this interval yields

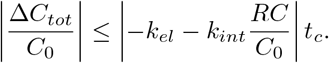

Substituting the expression for *t*_*c*_ and the bound on *RC* leads to the fQSSA validity condition (Fig 1d)

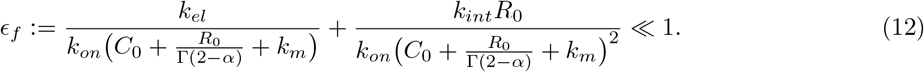

Setting *α* = 0 in Eq. 12 yields Γ(2 − *α*) = 1 and reduces *ϵ*_*f*_ exactly to the classical QSSA validity condition for the sTMDD model derived in our previous work [28], which is given by

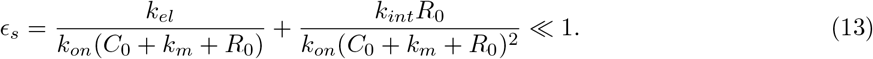

This consistency demonstrates that the fractional validity condition constitutes a consistent and natural generalization of the classical QSSA criterion in the presence of memory effects.

### Numerical method for the fTMDD and fQSSA models

The fTMDD model involves the modified Caputo operator 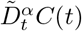, which requires a numerical approximation for efficient simulation. We discretize the fractional term using the Grünwald-Letnikov (G-L) scheme, a widely used and numerically stable approach for fractional-order models [23, 27, 29].

Let *t*_*n*_ = *nh* with a uniform step size *h >* 0 and denote *C*_*n*_ = *C*(*t*_*n*_). For 0 *< α <* 1, we approximate the Caputo derivative using a G-L convolution form, which is equivalent to the Riemann–Liouville representation supplemented with an explicit initial-condition correction under standard regularity assumptions:

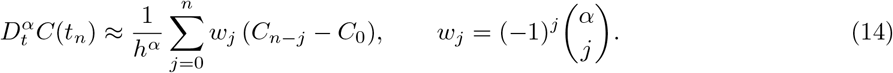

Using the modified Caputo operator (Eq. 2), the numerical approximation of 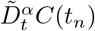 is therefore

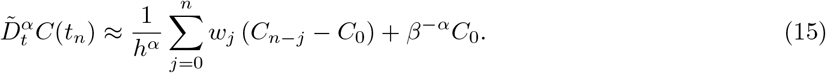

This discretization is consistent with the model definition and satisfies 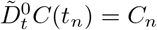, ensuring exact recovery of the sTMDD model in the limit *α* = 0.

At each time step, the fractional binding term is evaluated using Eq. 15 and the remaining ODE components are integrated using a fourth-order Runge-Kutta scheme. This hybrid method provides an efficient and stable numerical solver for the fTMDD model.

For the fQSSA model, the same discretization is used for the fractional derivative, while the free drug concentration is reconstructed at each time step by solving the algebraic QSSA constraint.

### Simulation design

To evaluate the dynamic behavior of the fTMDD model and assess the validity of the fQSSA across parameter regimes, we performed *in silico* simulations using the parameter sets summarized in Table 1. The simulations were designed to examine how memory effects and drug–target binding kinetics influence system dynamics and the accuracy of the reduced fQSSA model.

**Table 1.**
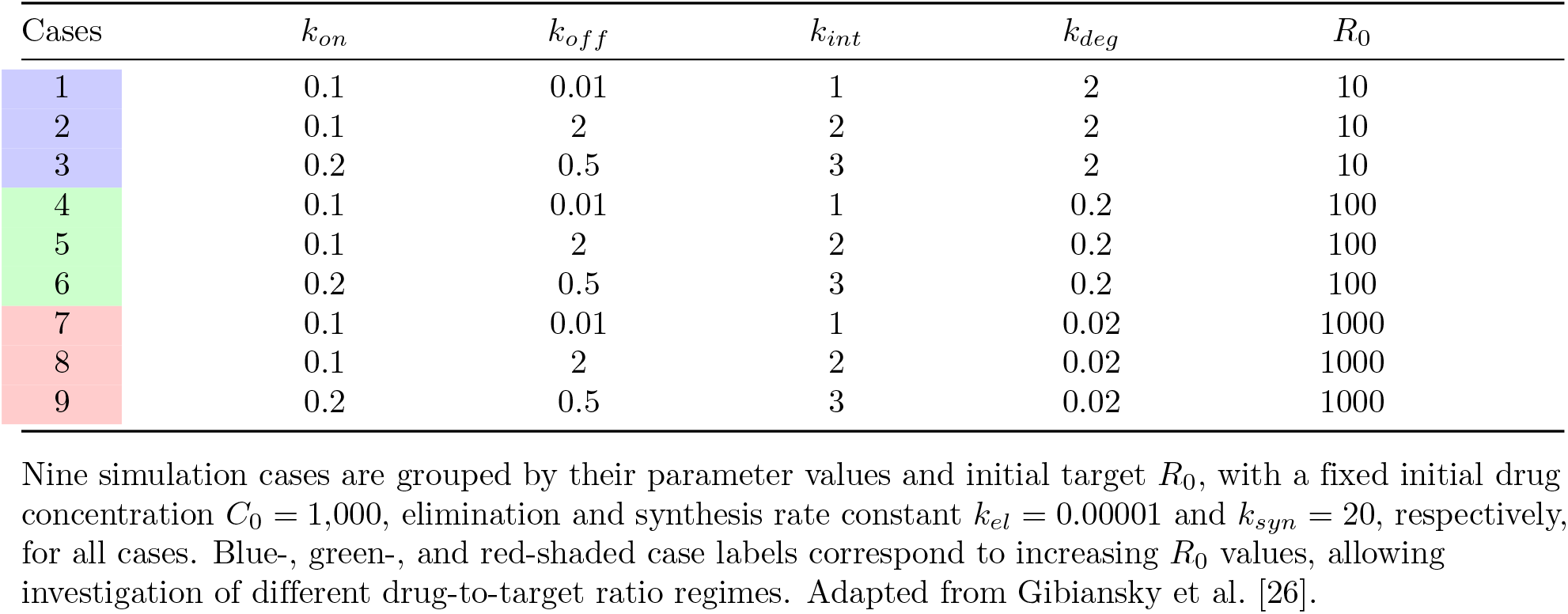
Parameter values used in the models.

| Cases | $k_{on}$ | $k_{off}$ | $k_{int}$ | $k_{deg}$ | $R_0$ |
| --- | --- | --- | --- | --- | --- |
| 1 | 0.1 | 0.01 | 1 | 2 | 10 |
| 2 | 0.1 | 2 | 2 | 2 | 10 |
| 3 | 0.2 | 0.5 | 3 | 2 | 10 |
| 4 | 0.1 | 0.01 | 1 | 0.2 | 100 |
| 5 | 0.1 | 2 | 2 | 0.2 | 100 |
| 6 | 0.2 | 0.5 | 3 | 0.2 | 100 |
| 7 | 0.1 | 0.01 | 1 | 0.02 | 1000 |
| 8 | 0.1 | 2 | 2 | 0.02 | 1000 |
| 9 | 0.2 | 0.5 | 3 | 0.02 | 1000 |

Nine simulation cases are grouped by their parameter values and initial target $R_0$ , with a fixed initial drug concentration $C_0 = 1,000$ , elimination and synthesis rate constant $k_{el} = 0.00001$ and $k_{syn} = 20$ , respectively, for all cases. Blue-, green-, and red-shaded case labels correspond to increasing $R_0$ values, allowing investigation of different drug-to-target ratio regimes. Adapted from Gibiansky et al. [26].

Nine parameter combinations were considered, grouped according to increasing initial target concentrations *R*_0_ (blue-, green-, and red-shaded cases in Table 1). Within each group, the initial drug concentration was fixed at *C*_0_ = 1000, while *R*_0_ was varied over one order of magnitude (*R*_0_ = 10, 100, 1000), enabling systematic exploration of different drug-to-target ratio regimes. For each *R*_0_, three combinations of (*k*_*on*_, *k*_*off*_, *k*_*int*_) were selected to represent distinct kinetic scenarios.

Parameter values were adapted from a previous TMDD study [26] and modified to span a broad range of physiologically relevant conditions, from target-excess (low *C*_0_*/R*_0_) to drug-saturated regimes (high *C*_0_*/R*_0_). This design enables a systematic assessment of how the memory parameter *α* and the ratio *C*_0_*/R*_0_ jointly influence the validity of the fQSSA model.

Using the numerical scheme described above, simulations were conducted for varying values of *α*. The resulting trajectories were used to compare the fTMDD and fQSSA models and to test whether the proposed validity condition *ϵ*_*f*_ ≪ 1 correctly predicts the parameter regimes in which the fQSSA provides an accurate approximation of the dynamics.

### Application: QSSA and fQSSA comparisons in recombinant human erythropoietin (rhEPO)

#### Data and study design

Plasma rhEPO concentration–time data were obtained from previously published studies in premature very low birth weight (VLBW) infants and normal adults following intravenous administration [30]. Three mean dose levels were considered in both populations: 10, 100, and 500 U/kg.

For each dose group, mean concentration–time profiles were used for model fitting. Sampling times (7-9 points) ranged from early post-dose (approximately 0.01 h) to 24 h, capturing both distribution and elimination phases. The observed concentrations span multiple orders of magnitude, motivating the use of log-scale error metrics in parameter estimation. Specifically, the QSSA and fQSSA models were implemented with an additional peripheral compartment to account for the distributional dynamics of rhEPO.

This extension is based on the fQSSA formulation derived above, with the modification that drug distribution between the central and peripheral compartments is explicitly incorporated. Accordingly, the model consists of three state variables: the central total drug concentration *C*_tot_(*t*), the peripheral drug concentration *C*_*p*_(*t*), and the total receptor concentration *R*_tot_(*t*). The free drug concentration *C*(*t*) is recovered through an algebraic quasi-steady-state relation.

The resulting system is given by

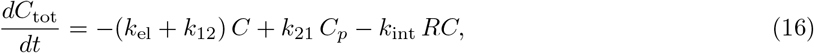

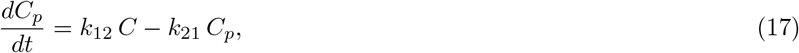

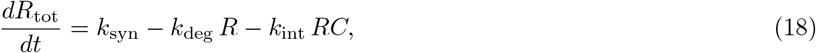

where

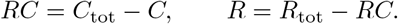

Under the QSSA, the free drug concentration *C* is determined by

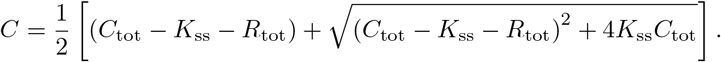

Under the fQSSA, the algebraic relation is replaced by

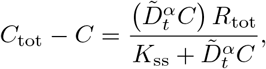

where 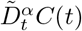 denotes the modified Caputo fractional derivative defined in Eq. (2). The initial conditions are specified as

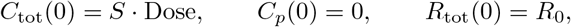

with *R*_0_ = *k*_syn_*/k*_deg_. Here, *S* is a dose-scaling parameter that converts the administered dose (U/kg) into an effective initial concentration in the central compartment. This parameter implicitly accounts for factors such as the apparent volume of distribution and unit conversion between dose and concentration. This formulation assumes that the administered dose instantaneously distributes into the central compartment, while the peripheral compartment is initially drug-free and the receptor system is at steady state prior to dosing.

#### Model fitting strategy

The QSSA and fQSSA models were fitted separately to infant and adult datasets. All dose groups within each population were fitted simultaneously to ensure parameter consistency across doses. Model simulations were initialized at *t* = 0 and evaluated at the experimental sampling times.

Model parameters were estimated by minimizing the log-scale discrepancy between observed and predicted drug concentrations:

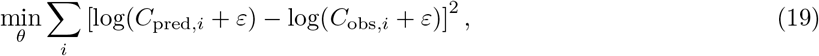

where *ε* is a small constant introduced to ensure numerical stability at low concentrations. Optimization was performed using a nonlinear least-squares algorithm (MATLAB `lsqnonlin`).

#### Model evaluation

Model performance was assessed using complementary criteria. Goodness-of-fit was evaluated via the residual sum of squares (RSS) and the log-scale root mean squared error (log-RMSE). Model complexity was accounted for using the corrected Akaike Information Criterion (AICc) and the Bayesian Information Criterion (BIC). Visual predictive performance was further assessed by comparing predicted concentration–time profiles with observed data across all dose groups, with particular attention to the early distribution and terminal elimination phases.

## Results

### As *α* increases, the system exhibits stronger deviation from classical Markovian dynamics

While the sTMDD model follows classical mass-action kinetics based solely on instantaneous drug concentration, the fTMDD model incorporates history-dependent effects through a Caputo fractional derivative. As a result, the contribution of past drug exposure influences current binding dynamics in the fTMDD model.

To characterize the impact of memory, we compared the temporal profiles of binding rates across different values of *α* (Fig 3). As *α* increases, the fTMDD model exhibits a slower decline compared to the sTMDD model, reflecting the increased influence of recent concentration changes and greater deviation from classical Markovian dynamics. This leads to prolonged drug retention and more persistent drug–target interactions.

**Fig 3.**
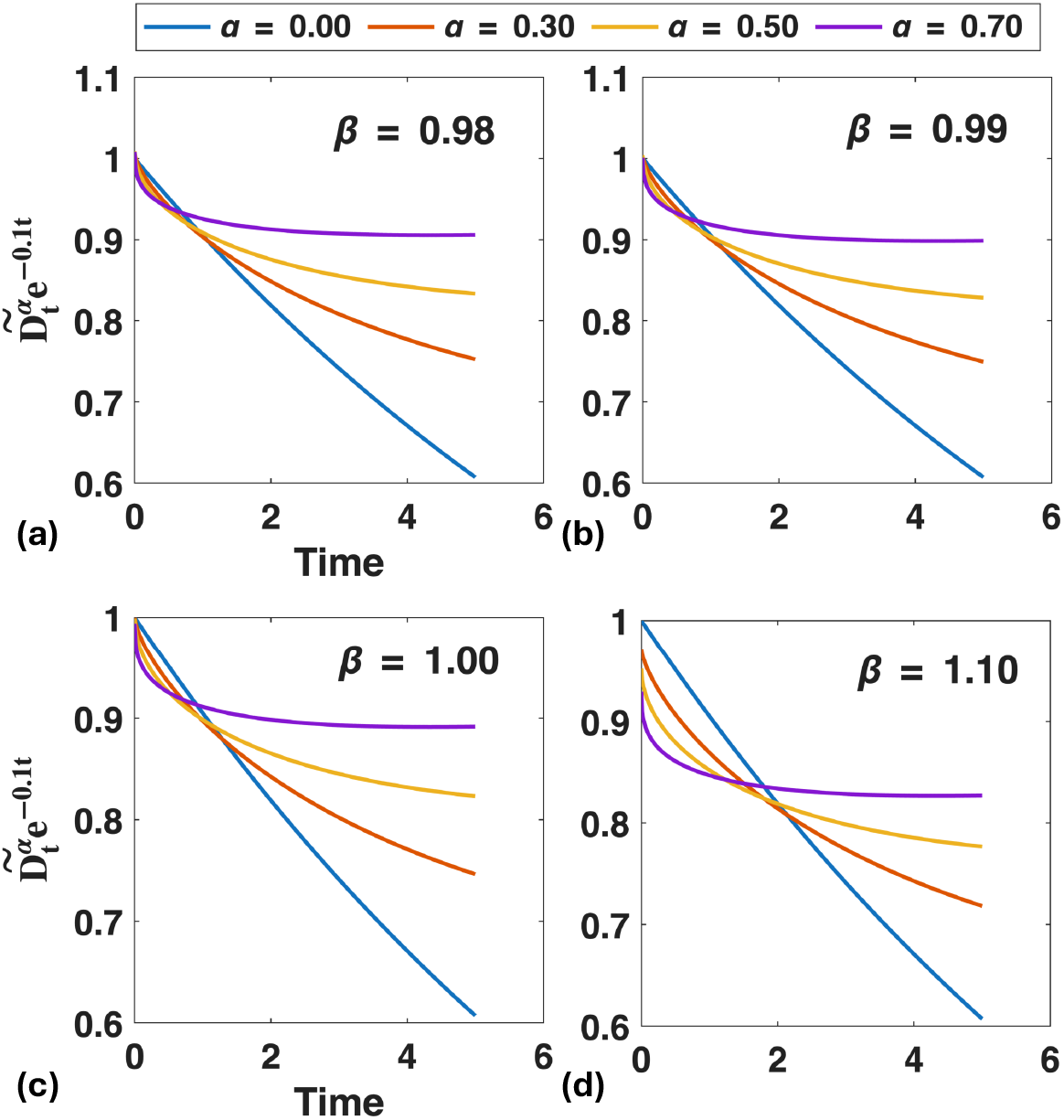
Influence of fractional order and memory scaling on fractional drug dynamics. Time evolution of the normalized fractional derivative term 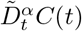 is shown for different fractional orders *α* = 0.00 (no memory), 0.30 (weak), 0.50 (mild), and 0.70 (strong), and memory parameters *β* = 0.98 (**a**), 0.99 (**b**), 1.00 (**c**), 1.10 (**d**). As *α* increases, the system increasingly deviates from classical Markovian dynamics, leading to responses that persist over longer timescales. Variations in *β* primarily affect the initial magnitude of the fractional term, while the qualitative influence of *α* on temporal behavior remains consistent across different *β* values.

The parameter *β* (Fig 3a–d) scales the initial contribution of the fractional memory term without affecting the qualitative dynamics determined by *α*. Accordingly, the memory scaling parameter *β* is fixed to *β* = 1 in all simulations to isolate and highlight the role of the fractional order *α*.

### fQSSA accurately captures memory-dependent pharmacokinetics

We next examined whether the fQSSA model can accurately approximate the fTMDD model across varying memory intensities (*α*), using the parameter set from Case 7 in Table 1. Both the fTMDD model (Fig 4a) and the fQSSA model (Fig 4b) consistently predict prolonged drug retention with increasing *α*, reflecting a slower decline of *C*_tot_(*t*) associated with changes in temporal weighting. The close agreement between the two models demonstrates that the fQSSA accurately captures the essential nonlinear and memory-dependent dynamics of the fTMDD system.

**Fig 4.**
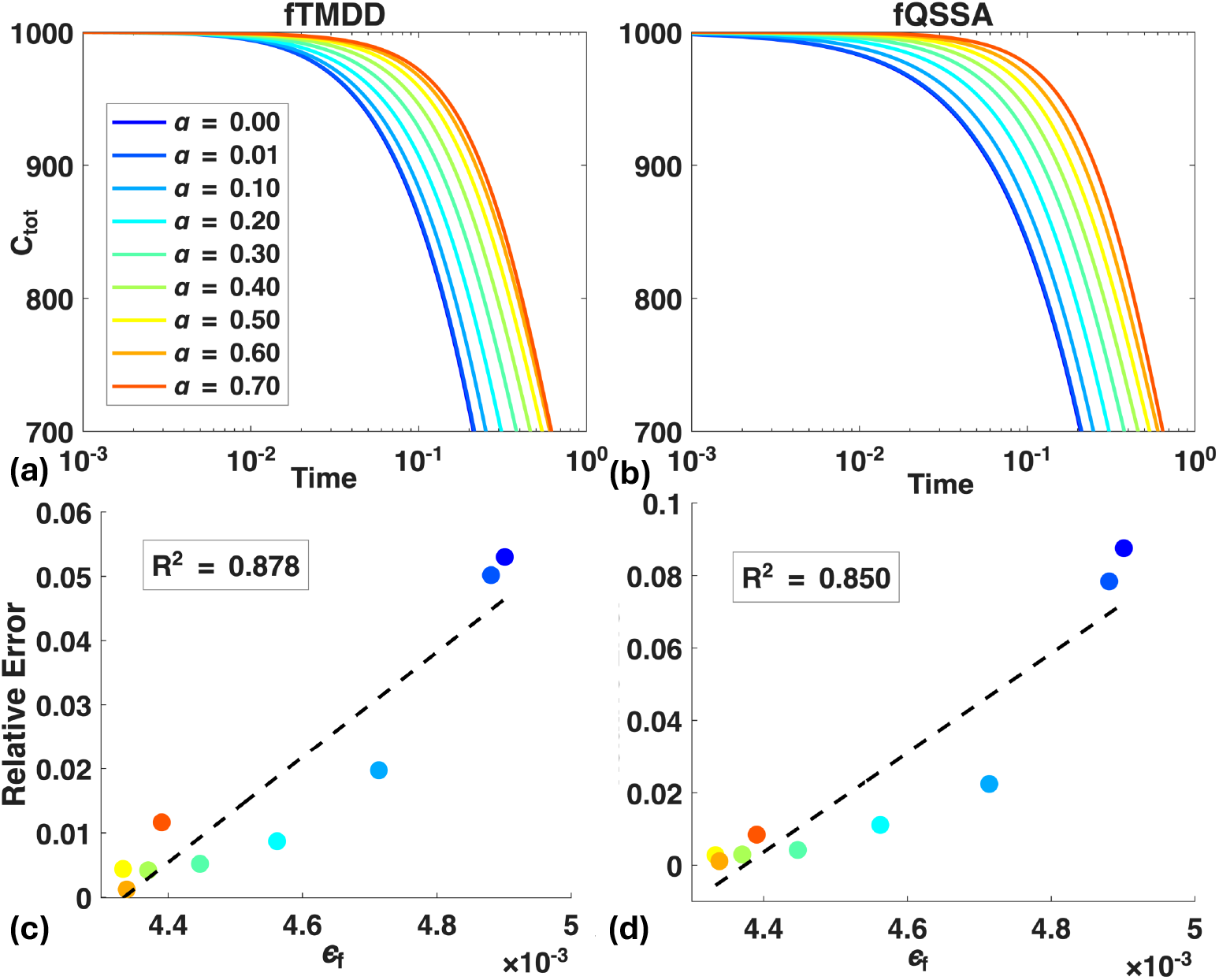
The fQSSA accurately approximates the fTMDD model across a wide range of memory parameters *α*. **(a–b)** Time evolution of total drug concentration *C*_tot_ under the fTMDD **(a)** and fQSSA (b) models. Both models exhibit prolonged drug retention as *α* increases, reflecting greater deviation from classical Markovian pharmacokinetic behavior. **(c–d)** Strong positive correlations are observed between the validity condition *ϵ*_*f*_ and the relative error at *t*_end_ = 1 **(c)** and *t*_end_ = 5 **(d)**. This indicates that *ϵ*_*f*_ reliably predicts QSSA accuracy across time.

Next, we evaluated the validity condition *ϵ*_*f*_ ≪ 1 (Eq. 12) by examining its relationship with the relative error in *C*_*tot*_(*t*), computed as the *L*^2^ norm over *t* ∈ [0, *t*_end_]. A strong positive correlation is observed between *ϵ*_*f*_ and the relative error (Figs 4c–d), with *R*^2^ = 0.878 for *t*_end_ = 1 and *R*^2^ = 0.850 for *t*_end_ = 5. These results confirm that smaller values of *ϵ*_*f*_ correspond to more accurate approximations, establishing *ϵ*_*f*_ as a reliable predictor of fQSSA accuracy.

### The drug-to-target ratio governs approximation accuracy of the fQSSA as predicted by the validity condition

Having established that the validity condition *ϵ*_*f*_ ≪ 1 reliably predicts the accuracy of the fQSSA, we next investigated the key determinant of *ϵ*_*f*_ : the initial drug-to-target ratio *C*_0_*/R*_0_. Our results show that increasing *C*_0_*/R*_0_ consistently reduces the validity indicator *ϵ*_*f*_, as shown in both the standard (*α* = 0) and fractional (*α* = 0.5) regimes (Figs 5a and 5c). This implies that greater drug availability leads to conditions that are more favorable for the QSSA approximation [28]. Indeed, the relative error between the fTMDD and QSSA models also decreases with increasing *C*_0_*/R*_0_ (Figs 5b and 5d). Together, these results confirm that *C*_0_*/R*_0_ governs the accuracy of the fQSSA, while *ϵ*_*f*_ provides a consistent quantitative predictor across memory conditions.

**Fig 5.**
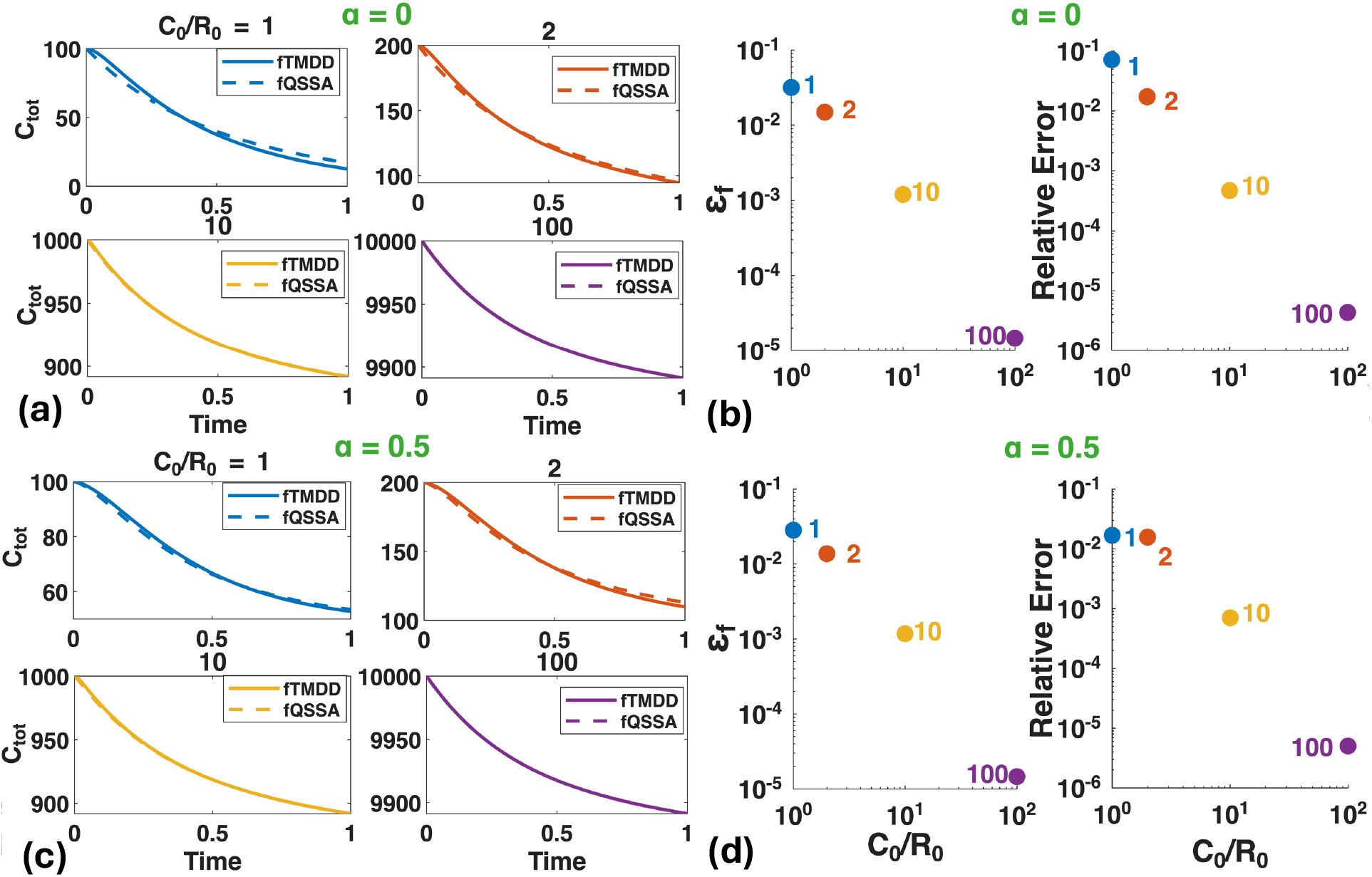
Drug-to-target ratio determines the accuracy of the fQSSA model as predicted by the validity condition. **(a–d)** Regardless of memory effects (**(a)** and **(b)** for *α* = 0; **(c)** and **(d)** for *α* = 0.5), the agreement between the fTMDD model and the fQSSA model improves as the initial drug-to-target ratio (*C*_0_*/R*_0_) increases (**(a)** and **(c)**). The relative error decreases with increasing *C*_0_*/R*_0_ and shows a strong positive correlation with *ϵ*_*f*_ (**(b)** and **(d)**).

The stronger dependence of *ϵ*_*f*_ on the drug-to-target ratio *C*_0_*/R*_0_, compared to *α*, can be understood from the structure of the validity condition *ϵ*_*f*_ (Eq. 12), which depends on the factor Γ(2 − *α*). Over *α* ∈ [0, 1), the variation of Γ(2 − *α*) is modest compared to the typically large variation in *C*_0_*/R*_0_. This indicates that memory effects act primarily as a secondary modulation, whereas the drug-to-target ratio remains the dominant control parameter.

### Systematic evaluation of fQSSA accuracy across drug-to-target ratios and memory intensities

To test whether the relationship between drug-to-target ratio, validity condition, and fQSSA accuracy remains robust across a wide range of parameter settings, we examined nine representative simulation cases (Table 1) with varying initial target concentrations (*R*_0_ = 10, 100, 1000) while fixing *C*_0_ = 1000, corresponding to drug-to-target ratios *C*_0_*/R*_0_ = 100, 10, 1. This allows us to systematically assess the accuracy of the fQSSA model across low to high saturation regimes under different memory intensities *α*.

Comparison of the fTMDD model (solid blue) and the fQSSA model (dashed red) shows a consistent pattern across all cases (Fig 6): the fQSSA model maintains high accuracy at large *C*_0_*/R*_0_ and deteriorates as the ratio decreases. This trend holds under both strong memory (*α* = 0.8) and weak memory (*α* = 0.1), indicating that the dominant role of *C*_0_*/R*_0_ persists across different memory intensities.

**Fig 6.**
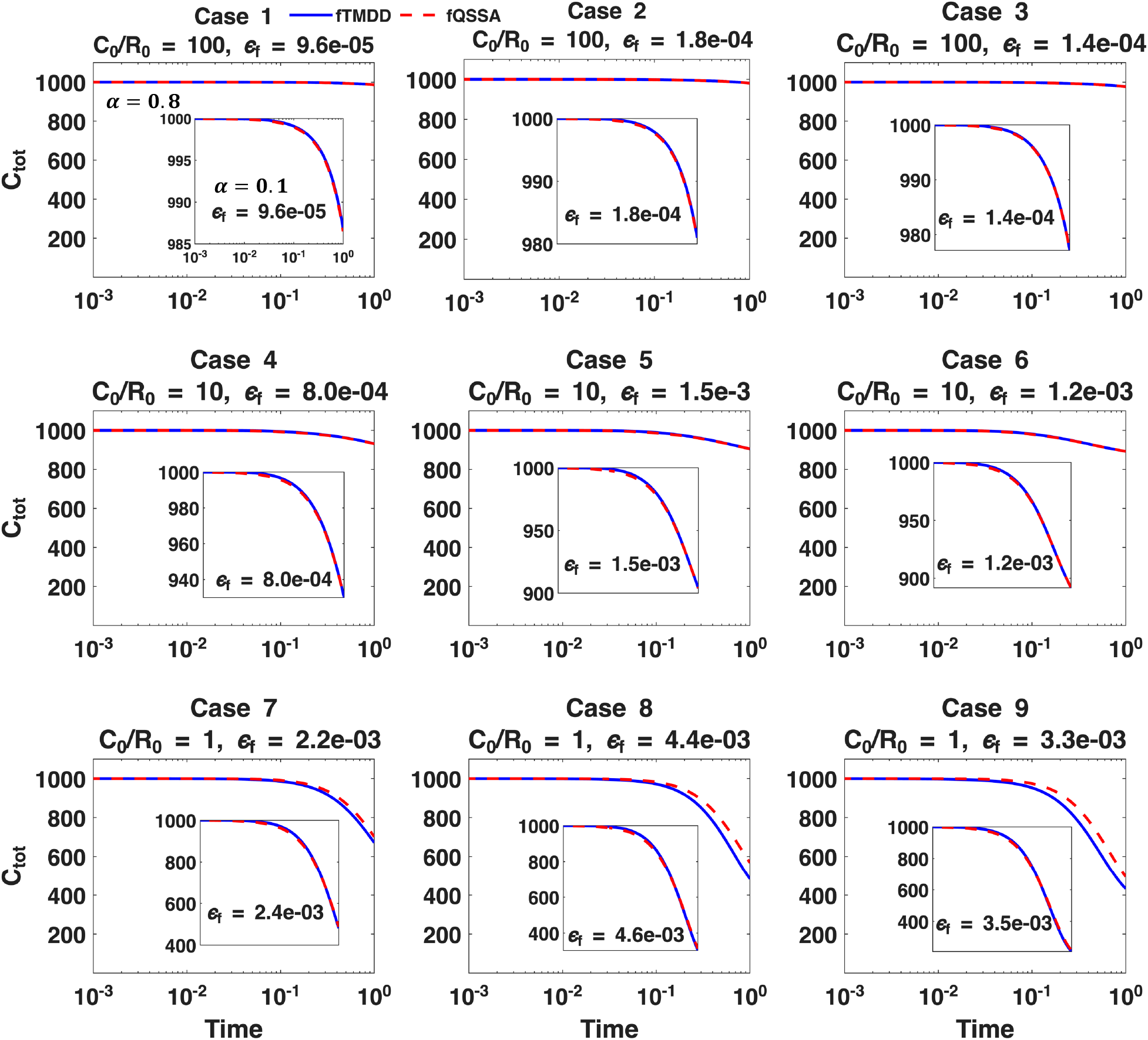
Comparison between the fTMDD model (solid blue) and the fQSSA approximation (dashed red) across nine different conditions. The main panels show results for memory intensity *α* = 0.8, while the insets show corresponding simulations with *α* = 0.1. Despite the difference in *α*, the approximation accuracy of the fQSSA shows little sensitivity to *α*, but is strongly influenced by the drug-to-target ratio (*C*_0_*/R*_0_ = 100, 10, and 1). Notably, Cases 7-9 with the lowest *C*_0_*/R*_0_ ratio exhibit the most visible relative error, which is consistently captured by the increasing values of the validity condition *ϵ*_*f*_ shown in each panel.

Importantly, larger relative errors consistently correspond to larger values of *ϵ*_*f*_, confirming that the validity condition remains predictive across parameter regimes. This relationship is further quantified by a strong correlation between relative error and *ϵ*_*f*_ in log–log scale (Figs 7a–b) for both *α* = 0.8 and *α* = 0.1.

**Fig 7.**
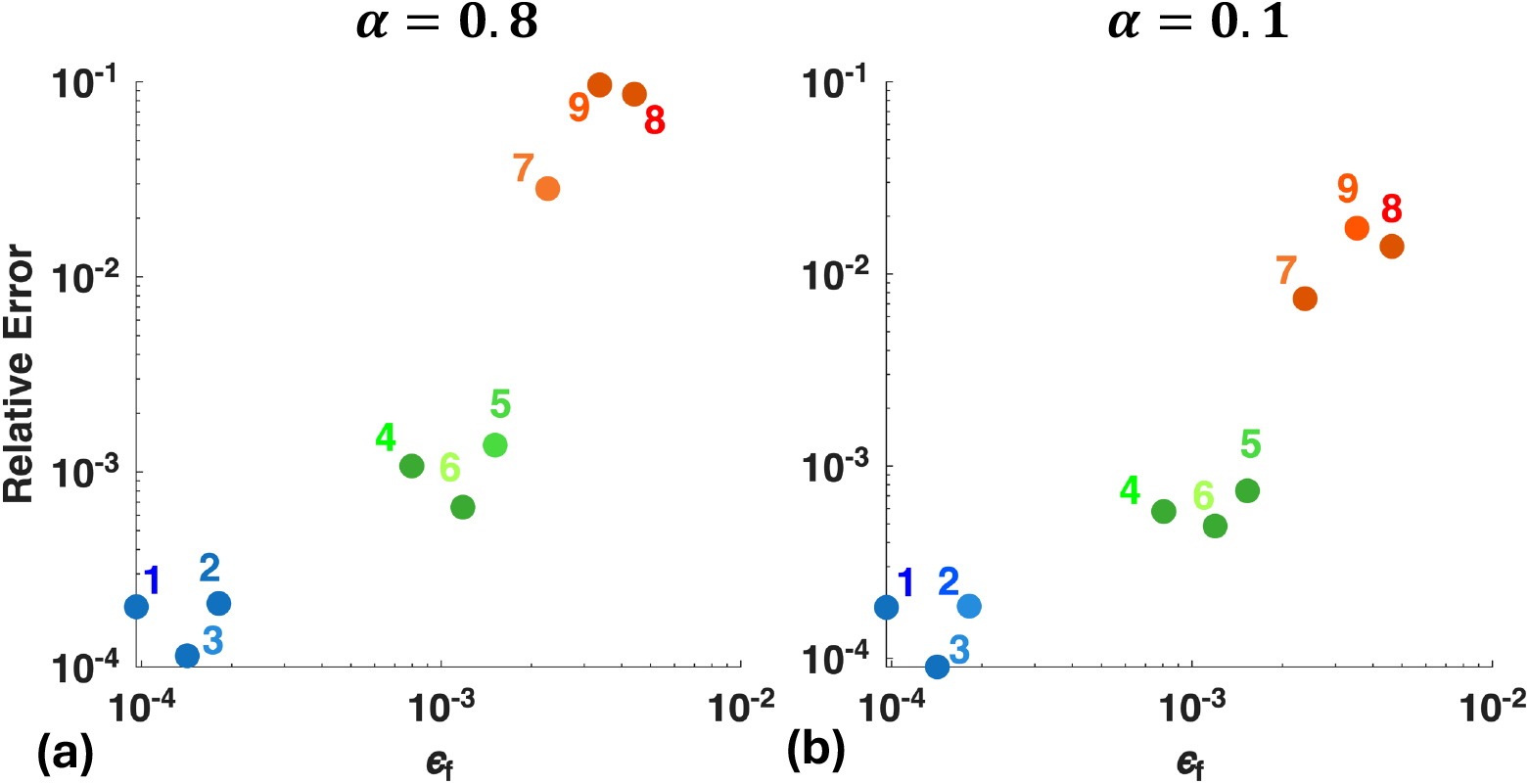
The relative errors and the validity condition of the fQSSA models in Fig 6. **(a–b)** The validity condition *ϵ*_*f*_ and the relative error are strongly correlated across nine different conditions for both *α* = 0.8 **(a)** and *α* = 0.1 **(b)**. This indicates that *ϵ*_*f*_ accurately predicts the accuracy of the fQSSA model, regardless of memory intensity.

Together, these results demonstrate that the interplay between drug-to-target ratio, validity condition, and fQSSA accuracy persists across a wide range of parameter settings. This highlights the generalizability and predictive power of the proposed validity condition *ϵ*_*f*_ for evaluating the accuracy of the fQSSA model.

### Comparison of QSSA and fQSSA model fits in application (rhEPO)

Having validated the fQSSA model through *in silico* simulations, we applied the proposed framework to plasma rhEPO concentration–time data in infants and adults. To assess model performance, we compared the QSSA [28] and fQSSA models.

Distinct differences between the two models are observed across the two populations (Table 2). In adults, the fQSSA model provides an improved fit compared to the QSSA model across all dose levels (10, 100, and 500 U/kg) (Fig 8a). The improvement is particularly evident in the early distribution phase and the terminal elimination phase, where the QSSA model deviates from the observed concentrations, while the fQSSA model captures these regions more accurately. This pattern is consistent across all dose groups. This improvement is also supported quantitatively. The log-RMSE decreases from 0.0872 (QSSA) to 0.0467 (fQSSA), and the AICc improves from −136.35 to −169.48, indicating a substantially better fit despite the increased model complexity.

**Table 2.** Estimated parameter values for QSSA and fQSSA models fitted to rhEPO data in infants and adults.

|  | Infants (QSSA) | Infants (fQSSA) | Adults (QSSA) | Adults (fQSSA) |
| --- | --- | --- | --- | --- |
| $k_{el}$ | 0.1754 | 0.1921 | 0.1038 | 0.0719 |
| $k_{12}$ | 0.3828 | 0.4492 | 0.6114 | 0.5017 |
| $k_{21}$ | 0.2853 | 0.3241 | 0.9433 | 3.0569 |
| $k_m$ | 82.2296 | 368.2743 | 297.4061 | 1752.0 |
| $k_{int}$ | 0.3976 | 0.1221 | 0.3168 | 1.5032 |
| $R_0$ | 87.4537 | 189.8036 | 214.4402 | 559.6741 |
| $k_{deg}$ | 0.1907 | 0.0484 | 0.0395 | 1.0e-05 |
| $k_{syn}$ | 16.6809 | 9.1772 | 8.4750 | 0.0056 |
| $S$ (Infants) | 14.1101 | 15.1228 | — | — |
| $S$ (Adults) | — | — | 7.5922 | 7.3916 |
| $\alpha$ | — | 0.0010 | — | 0.3320 |
| $\beta$ | — | 0.5173 | — | 1.0e+04 |

The fQSSA model additionally includes the fractional parameters $\alpha$ and $\beta$ . The steady-state constant $k_m$ is defined as $k_m = (k_{off} + k_{int})/k_{on}$ . $S$ denotes a proportional scaling factor relating model-predicted concentrations to observed data ( $C_{obs}(t) \approx S \cdot C_{pred}(t)$ ).

**Fig 8.**
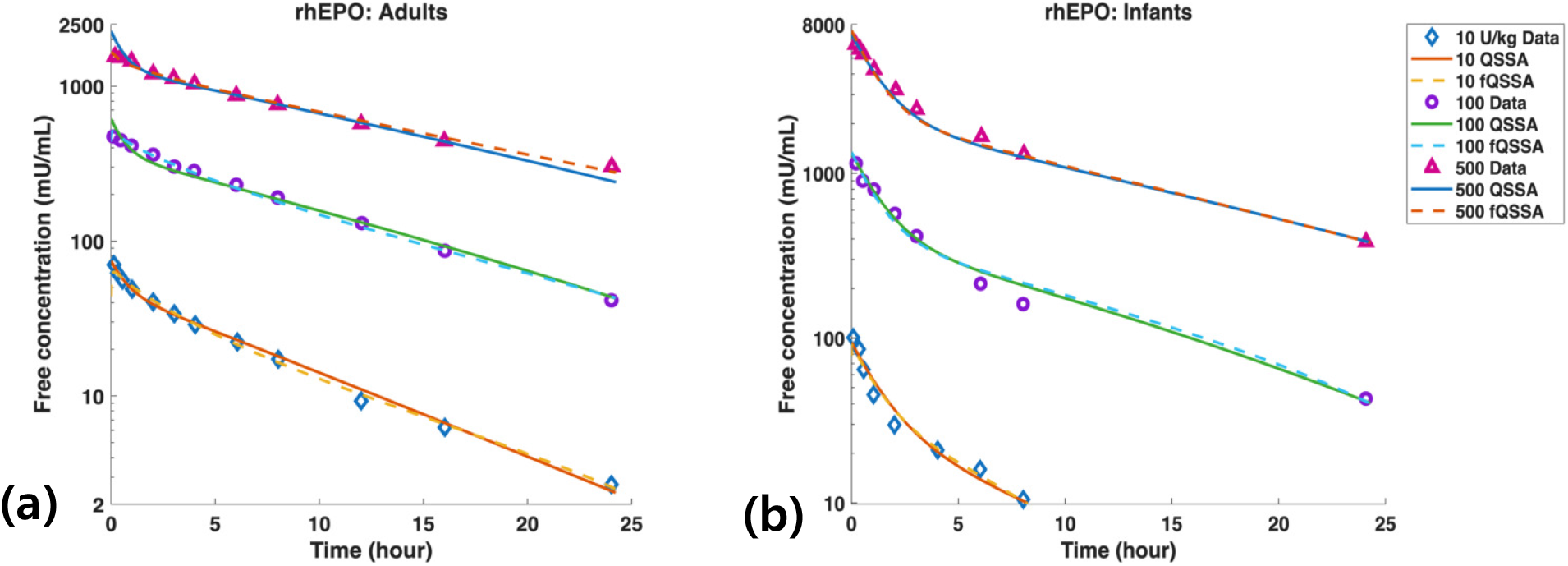
Comparison of QSSA and fQSSA models for plasma EPO concentration–time profiles. **(a–b)** Comparison of model fits in adults **(a)** and infants **(b)** following intravenous administration of 10, 100, and 500 U/kg. Symbols represent observed data, while solid and dashed lines denote QSSA and fQSSA model predictions, respectively. In adults, the fQSSA model improves the fit, particularly in the early and terminal phases. In infants, both models provide nearly identical fits, with the estimated fractional order *α* close to zero. Experimental data points were extracted from Fig 5 of D’Cunha et al. [30] using WebPlotDigitizer.

In contrast, in infants, both models yield nearly indistinguishable fits throughout the entire time course (Fig 8b). The predicted concentration profiles overlap across all dose levels, indicating that the fractional component does not provide a noticeable improvement. Consistently, the log-RMSE remains comparable (0.1097 vs 0.1167), while the AICc slightly worsens (−85.49 to −71.69), suggesting that the added model complexity is not justified for the infant data.

These population-level differences are consistent with the estimated fractional order. In infants, *α* ≈ 0.001, indicating negligible memory effects and near memoryless dynamics. In adults, *α* ≈ 0.33, together with improved model performance, indicates the presence of non-negligible memory effects, likely reflecting greater physiological heterogeneity in drug—target interactions.

Overall, this population-dependent contrast suggests that fractional dynamics reflect the degree of physiological heterogeneity of the system: adults exhibit history-dependent kinetics arising from greater biological complexity, whereas the relatively homogeneous pharmacokinetic environment in infants is adequately described by standard dynamics [31–33].

## Discussion

### The fTMDD model generalizes the sTMDD model by incorporating memory-dependent drug-target dynamics

The sTMDD models (such as Eq. 1) have played a central role in describing nonlinear drug dynamics arising from high-affinity drug–target interactions [1, 10]. These models assume that drug–target binding occurs instantaneously and depends solely on the current drug concentration. However, this assumption fails to capture physiological memory effects, such as prolonged receptor engagement or delayed conformational changes, which have been reported in several biologics and high-affinity drugs [34–36].

Recent studies have emphasized the need to account for temporal persistence in pharmacokinetics. Fractional-order models have been successfully used to describe anomalous diffusion, stochastic binding, and memory-dependent clearance processes [37–43]. These developments motivate a TMDD framework that incorporates history dependence while retaining a tractable mathematical structure.

In response, we propose an fTMDD model (Eq. 4) that incorporates memory effects via the Caputo fractional derivative. This operator captures the cumulative influence of past drug exposure through a power-law memory kernel, reflecting delayed or history-dependent drug–target interactions. The resulting formulation is governed by a single parameter, the fractional order *α*, which controls the strength of memory effects and provides a continuous bridge between memoryless dynamics (*α* = 0) and history-dependent behavior (0 *< α <* 1).

Importantly, when *α* = 0, the model reduces exactly to the classical sTMDD model (Eq. 1), ensuring backward compatibility and enabling direct comparison between memoryless and memory-dependent regimes within a unified framework. This property allows *α* to be estimated from data without imposing prior assumptions on the presence of memory effects.

Overall, the fTMDD model offers a unified and adaptive platform for describing nonlinear drug dynamics across both conventional and memory-aware regimes, while maintaining interpretability within established pharmacokinetic theory [44, 45].

### The fQSSA model bridges reduced complexity and biological realism in TMDD modeling

While the fTMDD model provides a flexible and mechanistically enriched framework for modeling memory-dependent drug-target kinetics, its expanded parameter space and computational demands can hinder parameter identifiability and practical applicability–especially when pharmacokinetic data are sparse [26, 46, 47]. To address this, we propose an fQSSA model (Eqs. 8–9) that reduces model complexity while preserving the essential nonlinear and memory-dependent dynamics.

QSSA has long served as a key model reduction strategy in biological systems, including PK–PD, particularly in systems where rapid drug-target binding or enzyme-substrate reactions permit timescale separation [48–54]. In sTMDD models, QSSA assumes that drug–target binding and complex internalization equilibrate rapidly compared to drug elimination [28, 55, 56]. It has also been widely applied to drug metabolism, enzyme-mediated clearance, receptor binding, and indirect response models [48, 51, 57–60]. However, these classical formulations assume memory-free dynamics and therefore cannot capture nonlocal temporal effects such as delayed drug responses, prolonged receptor engagement, or anomalous clearance behaviors observed in biologics and high-affinity systems.

The proposed fQSSA model extends this framework by incorporating fractional dynamics into the binding process, enabling reduced models to account for history-dependent effects. As demonstrated in our simulations (Figs 3–4), the fQSSA accurately reproduces the dynamics of the full fTMDD model across a wide range of memory intensities. This extension is further supported by the rhEPO application, where the fQSSA model captures population-dependent differences in drug dynamics beyond those described by the classical QSSA model. Together, these results establish the fQSSA as a theoretically consistent and practically useful generalization of QSSA for memory-affected systems.

### Validity condition *ϵ*_*f*_ determines the applicability of the fQSSA model

A key requirement for the practical use of the fQSSA model is to determine when it provides an accurate approximation of the full system. The applicability of QSSA under memory-dependent dynamics has remained largely unexplored, as previous studies have primarily focused on classical (Markovian) models [26, 28, 55].

To address this gap, we derived a validity condition *ϵ*_*f*_ (Eq. 12) that extends classical QSSA criteria to memory-dependent systems. This provides a systematic and theoretically grounded formulation of QSSA validity in fractional-order models.

The proposed metric *ϵ*_*f*_ quantifies the balance among binding and internalization kinetics, drug elimination, memory intensity *α*, and the initial drug-to-target ratio *C*_0_*/R*_0_. Importantly, it reduces to the classical QSSA validity condition *ϵ*_*s*_ (Eq. 13) when *α* = 0, ensuring consistency and backward compatibility with established theory [28].

To evaluate the predictive power of *ϵ*_*f*_, we compared it with the approximation error between the fTMDD and fQSSA models. Simulation results show that *ϵ*_*f*_ reliably predicts this error across a wide range of parameter regimes (Figs 6–7), with smaller values of *ϵ*_*f*_ consistently corresponding to higher accuracy. This relationship further reveals that the drug-to-target ratio *C*_0_*/R*_0_ is the dominant factor governing *ϵ*_*f*_, whereas the contribution of the fractional order *α* is comparatively modest.

Taken together, these findings establish *ϵ*_*f*_ as a theoretically grounded and practically useful criterion for determining when reduced-order modeling remains faithful to fTMDD dynamics. Because QSSA is widely used in PK–PD contexts—including drug metabolism, receptor binding, and enzyme-mediated processes—the proposed framework extends naturally to a broad class of fractional PK–PD models.

### Population-dependent role of fractional dynamics in rhEPO data

The rhEPO application provides a critical test of whether the fQSSA framework offers practical advantages beyond its theoretical appeal. Rather than simply confirming model improvement, we ask a more fundamental question: what does relevance-or irrelevance-of fractional dynamics reveal about the underlying pharmacokinetic system?

The answer proves to depend strongly on the population. In adults, the fQSSA model yields a markedly improved description of plasma rhEPO concentration–time profiles, particularly in the early distribution and terminal elimination phases (*α* ≈ 0.33, AICc: −136.35 → −169.48). This improvement suggests that drug–target interactions in adults are genuinely history-dependent, where the current binding rate is shaped not only by the present drug concentration but also by its past trajectory. From a mechanistic standpoint, such memory effects may arise from the increased biological complexity characteristic of the adult system, including variability in receptor expression, heterogeneous tissue distribution, and delayed intracellular trafficking [39, 42]. As organisms mature, changes in body size, organ structure, and tissue composition introduce multiple interacting timescales that classical Markovian models are fundamentally unable to capture. In this light, the fractional order *α* is not merely a fitting parameter but a physiologically meaningful quantity that reflects the degree of history dependence arising from biological complexity.

In infants, by contrast, the QSSA and fQSSA models yield nearly indistinguishable fits across all dose levels, with the estimated fractional order approaching zero (*α* ≈ 0.001). This indicates that the system is effectively memoryless, and that classical dynamics are sufficient to describe the underlying pharmacokinetic processes. Pharmacokinetic processes in neonates and infants are primarily governed by developmental physiology, including maturation of renal function and clearance pathways, which tend to produce more uniform and predictable drug disposition kinetics [32, 33, 61, 62]. In such a homogeneous setting, the time-scale separation inherent to the classical QSSA remains well-preserved, and fractional dynamics do not contribute meaningfully to model performance.

Taken together, these findings suggest that the fractional order *α* may serve as an indicator of physiological heterogeneity in drug—target interactions: when *α* ≈ 0, the system is well approximated by classical memoryless dynamics; as *α* increases, history-dependent kinetics become increasingly important and fractional models become indispensable. More broadly, this population-dependent behavior highlights that fractional models are particularly beneficial in systems where such heterogeneity and memory effects are pronounced, while simpler classical models remain sufficient in more homogeneous settings. This underscores the potential of the fTMDD and fQSSA frameworks as adaptive modeling tools capable of capturing both homogeneous and heterogeneous pharmacokinetic regimes within a unified formulation. In this sense, the fQSSA framework provides not only a modeling tool but also a data-driven mechanism for determining when memory effects should be incorporated in pharmacokinetic analysis.

### Current challenges and future directions in fTMDD modeling

While the fTMDD (Eq. 4) and fQSSA models provide a promising framework for modeling memory-dependent pharmacokinetics, several important challenges remain.

A primary issue is the interpretation and estimation of the fractional order *α*. Although *α* serves as a compact parameter quantifying memory intensity in drug–target interactions, its direct physiological meaning remains unclear. Existing studies support the presence of memory effects in pharmacokinetics [35, 37], but further experimental work is required to link *α* to specific mechanisms such as receptor recycling, delayed binding, or anomalous transport processes [42, 63–66].

Closely related to this issue is the challenge of estimating *α* from empirical data. Because *α* characterizes long-range memory through a power-law kernel, its estimation can be viewed as a kernel identification problem. Promising approaches include Bayesian inference and physics-informed neural networks, although their application to pharmacokinetic systems remains an open area of research [67–70].

Beyond parameter estimation, extending the current framework to more complex settings represents an important direction for future work. In particular, the validity of QSSA is known to differ between deterministic and stochastic systems [71–74], as well as in spatially heterogeneous environments [75, 76]. Deriving corresponding validity conditions for fractional models in these contexts will be essential for accurately capturing noise-driven and spatially structured pharmacokinetic dynamics.

In addition, most therapeutic settings involve multi-drug interactions or combination therapies that can alter binding kinetics, target availability, and elimination pathways [12, 13]. Recent studies have begun to extend TMDD models to competitive multi-drug systems, highlighting the importance of modeling interactions among multiple agents sharing common targets [77]. Extending the fTMDD and fQSSA frameworks to incorporate such multi-species interactions, together with memory-dependent dynamics, represents a key next step toward realistic pharmacokinetic modeling.

Despite these challenges, this work establishes a systematic and extensible foundation for incorporating memory effects into TMDD modeling. By linking classical and fractional formulations through a unified framework, and by introducing both reduced models and an explicit validity condition, we provide a principled approach for analyzing nonlinear pharmacokinetic systems with memory. Although developed for TMDD, the proposed methodology is broadly applicable to a wide range of fractional PK–PD models, including drug metabolism, receptor binding, and indirect response systems [20, 42, 54, 78–80].

## Supporting information

**S1 Table. Goodness-of-fit statistics for QSSA and fQSSA models fitted to rhEPO data in infants and adults**. Lower values indicate better fit for all metrics.

**S1 Data. Experimental data points extracted from previous publications [30] used for application**.

**S1 Code. MATLAB scripts implementing the standard TMDD, fractional TMDD and QSSA models, including simulation, parameter estimation, and figure reproduction used in this study**.

